# Strongyle-resistant sheep express their potential across environments and leave limited scope for parasite plasticity

**DOI:** 10.1101/2020.06.19.161729

**Authors:** G. Sallé, V. Deiss, C. Marquis, G. Tosser-Klopp, J. Cortet, D. Serreau, C. Koch, D. Marcon, F. Bouvier, P. Jacquiet, N. Holroyd, A. Blanchard, J.A. Cotton, M.M. Mialon, C. Moreno-Romieux

## Abstract

**Introduction:** Drug-resistant parasites threaten livestock production. Breeding more resistant hosts could be a sustainable control strategy. Environmental variation may however alter the expression of genetic potential and directional selection toward host resistance could initiate an arms race between the host and its parasites.

**Methods and Results:** We created sheep lines with high or low resistance to *Haemonchus contortus*. We first exposed both lines to chronic stress or to the infection by another parasite *Trichostrongylus colubriformis*, to test for genotype-by-environment and genotype-by-parasite species interactions respectively. Overall, between-line divergence remained significant across environmental perturbations. But we found that the impact of chronic stress on *H. contortus* infection varied among families and that divergence was reduced during infection by *T. colubriformis*. Second, we quantified genomic and transcriptomic differences in *H. contortus* worms collected from both lines to identify components of an arms race. We found no evidence of genetic differentiation between worms from each line. But survival to more resistant hosts was associated with enhanced expression of cuticle collagen coding genes.

**Discussion:** Breeding for resistance hence remains a sustainable strategy that requires to anticipate the effects of environmental perturbations and to monitor worm populations.

## Introduction

Gastro-intestinal nematodes (GIN) impose a significant burden to human health and livestock worldwide. Repeated systematic anthelmintic drug treatments have favoured the rapid selection of drug-resistant isolates across continents (Kaplan & Vidyashankar, 2012), rendering sheep farming impossible in some cases (Blake & Coles, 2007). Concerns about environmental side-effects associated with anthelmintic drug metabolites (Verdú et al., 2018) have also driven research effort to develop alternative control strategies.

Breeding more resistant individuals is a promising alternative. Indeed, domestic (Bishop, 2012; Gruner, Bouix, & Brunel, 2004; Woolaston, 1992) and wild populations (Gold et al., 2019; Smith, Wilson, Pilkington, & Pemberton, 1999; Sparks et al., 2019) often show heritable genetic variation for resistance to parasite infection that breeding programs could exploit. This is despite the theoretical predictions that would expect the rapid fixation of favorable alleles through positive selection (Kruuk, Slate, & Wilson, 2008).

Several factors have been proposed to explain this discrepancy (Kruuk et al., 2008; Lazzaro & Little, 2009; Seppälä, 2015; Seppälä & Jokela, 2016). First, sheep resistance to GIN infection has a polygenic architecture. This has been described with genome-wide resolution in commercial sheep populations (Kemper et al., 2011) and evidence of multiple Quantitative Trait Loci (QTL) with mild effects were found across European breeds (Riggio et al., 2014; Sallé et al., 2012, 2014) or in sheep lines bred for divergent susceptibility toward GIN infection (McRae, McEwan, Dodds, & Gemmell, 2014). This genetic network likely causes functional trade-offs between immune response and fitness as a result of pleiotropy, although weak positive (Assenza et al., 2014; Bishop, Bairden, McKellar, Park, & Stear, 1996; Bouix et al., 1998) or negative (Douch, Green, Morris, & Hickey, 1995; Eady et al., 1998) genetic correlations between resistance and growth traits were found in domestic populations.

Second, environmental perturbations likely contribute to maintaining genetic variation in the host population by allowing different genotypes to express maximal fitness across conditions, as a result from host genotype x environment (*G_h_ x E*) interaction (Hoffmann & Merila, 1999; Lazzaro & Little, 2009; Lynch & Walsh, 1998; Seppälä & Jokela, 2016) or by inflecting the strength and direction of selection (Hayward et al., 2018). For instance, analyses of Faecal Egg Count (FEC) data from commercial Merino sheep revealed increased heritability under the lowest and the highest parasite exposure, as a result from variation in sire estimated breeding values across environments (Hollema, Bijma, & van der Werf, 2018; Pollott & Greeff, 2004).

Third, GIN can promote survival of less common host genotypes that they are less prone to invade (host genotype x GIN genotype interaction, *G_h_ x G_p_*), thereby fostering an adaptive arms race with their hosts (Van Valen, 1973). Strong directional selection toward higher resistance in the host population could hence disrupt the host-GIN coevolutionary dynamics. While current knowledge suggests that parasites do not adapt to resistant sheep (Kemper, Elwin, Bishop, Goddard, & Woolaston, 2009), contradictory observations from *Heligmosomoides polygyrus* infected mice suggest alterations in both parasite fecundity and immunomodulatory ability can develop after repeated infection (Lippens, Faivre, & Sorci, 2017). In addition, GIN populations like *H. contortus* have ample genetic diversity for selection to act upon (Sallé et al., 2019), and they also demonstrate enough transcriptional plasticity to circumvent the immune response of their hosts (Sallé et al., 2018). It is hence unclear how strong directional selection toward resistance to a single GIN species would affect that species and how this selection could promote rewiring of GIN species assemblage by reducing resistance to other parasite species. Limited evidence from domestic Merino (Woolaston, Barger, & Piper, 1990) or Romane sheep breeds (Gruner et al., 2004) however suggest that selection for resistance to *H. contortus* confers significant but incomplete cross-resistance to *Trichostrongylus colubriformis*, an intestinal GIN.

Therefore, abrupt variation in environmental conditions could release host cryptic variation overlooked under controlled conditions, and directional selection for host resistance could disrupt host-parasite interactions. This is critical to the sustainability of the breeding option for the control of GIN populations. To investigate that matter, we created divergent sheep lines selected for resistance or susceptibility to *H. contortus* infection. We monitored their resistance potential under chronic stress or against *T. colubriformis* infection, and we measured genomic and transcriptomic modifications occurring between *H. contortus* worms from both sheep backgrounds. Our observations suggest that limited but significant *G_h_* x *E* and *G_h_* x *G_p_* interactions occur across considered environments, and that plastic expression of cuticular collagen may help the parasite to circumvent the immune response of its host.

## Materials and methods

### Creation of two divergent lines of sheep

A full description of the selection scheme has been provided in the supplementary technical note. Briefly, a QTL detection scan across European breeds identified eight regions significantly associated with resistance to GIN (Riggio et al., 2014). SNP from the 800K SNP chip located within these QTL regions (approximately 110 SNPs per region) were subsequently selected and genotyped using the KASP™ assay (LGC Genomics Ltd, UK) that consists in a competitive allele-specific PCR (He, Holme, & Anthony, 2014). A nucleus Romane flock (ancestral generation G0, n = 277) was genotyped for these markers and phenotyped for resistance to *H. contortus* by two consecutive challenges with 10,000 larvae, as previously outlined (Sallé et al., 2012).

A marker-assisted selection approach was subsequently applied to retain the most resistant and most susceptible G0 sires and ewes for conditional mating using single-step GBLUP (Aguilar et al., 2010; Christensen & Lund, 2010). Instead of relying only on the genomic relationship matrix (VanRaden, 2008; Yang et al., 2010), this approach models the phenotype as the sum of fixed effects and a random animal effect estimated from a blended relationship matrix ***H*** that accounts for differences between pedigree and genomic information (Aguilar et al., 2010; Christensen & Lund, 2010). The relative weight given to the pedigree or genomic information is defined by a scaling parameter that ranges from 0 (pedigree only relationship matrix) to 1 (marker only relationship matrix) and was, in our case, set to 0.5 (Aguilar et al., 2010). The genomic relationship matrix used here consisted of the raw genomic information scaled by the parameter 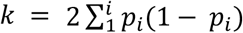 where *p_i_* refers to SNP *i* allele frequency (this aims to make the genomic relationship matrix similar to the pedigree relationship matrix). Matrix was then weighed to facilitate matrix inversion (VanRaden, 2008).

Using these genomic estimated breeding values (***geBVs***), the six most extreme G0 sires (three at each end of the geBV distribution) were mated with 55 and 63 resistant and susceptible G0 ewes respectively (among 118 females with breeding value estimations), to create 236 lambs (generation G1).

Out of these 236 G1 lambs, a subset of 180 lambs were selected for genotyping with the 1000-SNP chip, according to their expected breeding value (average of their parent breeding values, ***eBV***). eBVs for first and second infection FEC were computed using a model including known fixed effects (litter size, sex) and an individual random effect estimated from the pedigree relationship matrix. Their geBVs were subsequently derived using their genotype information and the SNP effect calculated in G0. 87 G1 lambs were retained for the experiment.

A second generation of lambs was produced following the mating of six G1 rams (three within each line, selected on their proper breeding values) with 82 ewes selected among G1 ewes (n = 23 and 19 selected on their eBVs for both R and S lines respectively) and G0 ewes (n = 19 and n = 20 selected on their eBVs for both R and S lines respectively). A total of 111 lambs were created in generation G2 (from 55 ewes), out of which 80 lambs were subsequently selected within each line according to their eBVs. Analyses were run with AIREMLF90 and BLUPF90 for eBVs and geBVs estimations respectively (Misztal et al., 2002).

### Behavioural treatment to establish how sheep resistance potential holds in stressful environments

Animal experiments and experimental procedures were approved by the French Ministry for Higher Education and Research and the Centre Val de Loire ethics committee under agreement numbers 2015010811379451_v4 and APAFIS#8973-2017022108587640_v3. Behavioural treatment was applied to 84 enrolled G1 lambs allocated to four indoor pens, each housing 22 females or 20 males with equal proportion of lambs from both lines. Half of the sheep were submitted to a stress treatment or a control treatment seven weeks prior to the first *H. contortus* infection. This behavioural treatment lasted throughout the experimental infection. The control treatment involved mild enrichment: sheep had access to a wool brush and were exposed to daily positive tactile contacts with humans. Twice a week, a familiar sheep keeper entered the pen, stayed passive and gave positive contacts to sheep that initiated contacts. The chronic stress treatment consisted in moving animals from their free-range pens to individual cages where they remained locked down and isolated from their mates once a week. Isolation was applied until the end of the second artificial challenge and lasted 20 minutes in the first week, 40 min the following month and 10 min afterwards.

Measurements focused on behavioural reactivity and standing-lying behaviour that were recorded before the onset of behavioural treatments, five weeks later, and 14 days after the first and second infections. Additional recordings of standing-lying behaviour were performed on the same day and just before experimental infection took place.

Behavioural reactivity was measured by an arena test (2 m x 7 m dirt floor with 2 m high solid walls and 7 equal-sized areas delimited by lines on the floor) that evaluate social attractiveness of two lambs for their flock-mates (n = 3, kept behind a wire mesh) or for a standing human being. The first phase of the test (1 min) evaluated reaction to novelty and social isolation (measured by locomotion, *i.e*. number of squares entered, and vigilance, *i.e*. head in upright position and ears perpendicular to the head). Tested lambs were first isolated from their flock-mates (hidden behind a curtain) for 1 min before the curtain was lifted to allow social proximity between tested lambs and their mates. The second phase consisted in an operator entering the arena and standing in front of the wire mesh for 1 min to measure lamb social attraction and reaction to a stationary person (locomotion, vigilance, bleat, physical contact with their flock-mates and with the operator). Lamb behaviour was recorded with a camera (Sony SPT-MC128CE, Sony Corp., Tokyo, Japan) and video recorder (Sony SVT-1000P, Sony Corp., Tokyo, Japan). Vocalisations were recorded directly by an observer hidden from lambs.

Standing-lying behaviour was recorded on females with an accelerometer (HOBO^®^ Pendant^®^) attached with a cohesive bandage to the lateral side of the hind leg (left or right in half the lambs each). The accelerometer was positioned so that the x-axis was vertical and towards the ground, and the y-axis parallel to the ground and towards the rear of the animal.

### Artificial infection with *H. contortus* and *T. colubriformis*

Lambs were kept indoors during the whole experiments to prevent natural GIN infection. The G1 lambs were challenged twice with 10,000 *H. contortus* infective larvae, given orally after three months of age. At the end of the first infection (30 days post-infection, dpi), lambs were drenched with ivermectin (2.5 ml/10 Kg body weight of Oramec^®^, Merial, France) and left for a resting period of two weeks before another infection took place with the same infection dose. Lambs were weighed on the same day and Faecal Egg Counts were quantified at 24- and 30-dpi.

To evaluate whether genetic resistance was sustained in the face of an intestinal GIN, the second generation of sheep from both divergent lines were either infected by *H. contortus*, or by 10,000 infective *T. colubriformis* larvae. In the latter case, FEC were measured at 24, 30 dpi after the first infection and at 30 dpi after the second infection. Lambs were weighed before and at 14 and 30-days post infection.

### Transcriptomic profiling of *H. contortus* communities from resistant and susceptible sheep

Pools of 50 male worms were collected from four susceptible and four resistant G1 sheep and snap-frozen in liquid nitrogen. DNA and RNA were simultaneously extracted using the AllPrep DNA/RNA/miRNA Universal Kit^®^ (Qiagen, UK) following manufacturer’s instructions on two pools of 10 worms and an additional pool of 30 worms, yielding three replicates per sheep. RNAs were sequenced on two lanes of HiSeq v4 using 75-bp paired-end reads. Adapter-trimmed reads were mapped onto the *H. contortus* genome v4 (Doyle et al., 2020) using STAR v2.5 (Dobin et al., 2013). Read counts were subsequently normalized for length and GC content biases using the cqn package v1.28. Differentially expressed genes between worms from resistant and susceptible sheep were identified by DESeq2 v1.22.1 (Love, Huber, & Anders, 2014) and VOOM (Law, Chen, Shi, & Smyth, 2014) as implemented in limma v3.38.3. Genes with absolute fold-change higher than 1 and with Bonferroni-Hochberg (Benjamini & Hochberg, 1995) adjusted *P* evalues below 5% were regarded as significantly differentially expressed. To avoid relying on a single read mapping pipeline, pseudo-alignment was also implemented with Salmon v0.11.3 (Patro, Duggal, Love, Irizarry, & Kingsford, 2017) correcting for GC content, position-specific and sequence-specific biases using dedicated options (--gcBias, --seqBias, --posBias respectively). Quantification files were imported using the tximport package v1.10 (Soneson, Love, & Robinson, 2015) before identification of differentially expressed genes as described above. The intersecting set of differentially expressed genes between the three pipelines were retained as significant. Fold-changes and associated *P*-values estimated from STAR counts with the VOOM frameworks were reported, as estimates in this case were in general more conservative. Genomic positions, protein domains and predicted orthologs of differentially expressed genes in *Caenorhabditis elegans* were retrieved from WormBaseParasite v13.

Validation of the differentially expressed genes was run on independent pools of worms collected from three resistant and three susceptible G2 lambs. For each sheep genetic background, RNA was extracted from 6 pools of 50 worms using an RNeasy^®^ kit (Macherey-Nagel). Total RNA was used for oligo(dT) cDNA synthesis (SuperScript^®^ III First-Strand Synthesis System, ThermoFisher, France, 18080051). The resulting cDNA was diluted 1:250 for quantitative PCR and 1 *μ*l added to each reaction. qRT-PCR was carried out following the iQ SYBR GREEN supermix^®^ protocol (Biorad, France, 1708882) and was run in triplicate. Primers were designed using the Primer3Plus website and blasted against *H. contortus* genome (Doyle et al., 2020) using the WormBase Parasite interface (Howe, Bolt, Shafie, Kersey, & Berriman, 2017). Designed primers have been listed in supplementary Table 1. To identify genes with statistically different expression levels, measured Ct cycles were regressed upon gene, treatment group and their interaction, fitting house-keeping gene average Ct as a covariate and considering each biological replicate (n = 6 per gene) as a random effect. This mixed model was implemented with the R *nlme* package v.3.1-142 (Pinheiro, Bates, Debroy, & Sarkar, 2019). One sample (S-1) was an outlier and was removed from the analysis. Results were given as expression fold-change using the ΔΔCt method (Vandesompele et al., 2002).

### Genetic diversity of *H. contortus* communities from resistant and susceptible sheep

Genomic data were mapped onto the latest *H. contortus* reference genome using smalt v 0.7.4 (ftp://ftp.sanger.ac.uk/pub/resources/software/smalt/smalt-manual-0.7.4.pdf) with 90% identity for alignment and a maximum insert size of 2000 bp (-y 0.9 -i 2000 options respectively) and default k-mer length of 13. Bam files corresponding to worms collected from the same sheep were merged using the picard v2.14.0 MergeSamFiles tool. The resulting bam file was filtered further with samtools v0.1.19-44428cd to remove reads of poor quality (-q 20), with poor mapping quality (-Q 30), or unmapped (-F 0×4 -F 0×8) and to only retain properly mapped pairs (-f 0×2). Duplicate reads were removed using the Picard v2.14.0 REMOVE_DUPLICATE tool. The popoolation2 framework (Kofler, Pandey, & Schlotterer, 2011, p. 2) was subsequently applied to create a sync file used to compute FST in 10-Kbp wide sliding windows (step of 1000 bp). Briefly, an mpileup file was built using samtools and used as an input to create a sync file using the mpileup2sync.jar tool. Indels were identified and removed from the sync file using the *identify-indel-regions.pl* and *filter-sync-by-gtf.pl* tools respectively. Windows with less than 100 SNPs, less than 50% of its width being covered or showing depth of coverage above 120X were discarded. Pairwise F_ST_ comparisons were averaged within- or between-groups. Regions showing outlying genetic differentiation (3 standard deviations from mean FST) between worms from resistant and susceptible sheep but not between worms from the same sheep lines (within-group) were regarded as putative candidates promoting survival in resistant sheep.

### Statistical analyses

Statistical analyses were implemented with R v3.5 unless stated otherwise (R Core Team, 2016). For every analysis, raw FEC data were normalized by a fourth-root transformation that outperformed the logarithmic transformation (Shapiro-Wilk’s test ranging between 0.96 and 0.90 for fourth-root transformed FEC but 0.90 and 0.84 for log-transformed FEC). Summary statistics of considered variables and detailed modeling outputs have been provided in supplementary tables 2 and 3 respectively. Considered response variables were either average FEC at first or second infection (average between measures taken at 24 and 30 days post-infection) or across the two infections (average across infections).

### Response to selection

To estimate phenotypic divergence in FEC between sheep lines, individual *H. contortus* infection data were pre-corrected for fixed environmental effects, *i.e*. sex, litter size, and generation (accounting for year effect), and an individual random effect, using the nlme package v3.1-140 (Pinheiro et al., 2019). Individual random effects were standardized to unselected G0 mean and standard deviation.

Responses to selection for FEC at first, second infection or across infections were evaluated within each generation by regressing individual random effects from *H. contortus* infected offspring upon their respective midparent values, computed as the average value of each lamb’s sire and dam (Falconer & Mackay, 1996). This regression coefficient provides an estimate for realized heritability (Falconer & Mackay, 1996) and was used to establish the asymmetry of response between resistant and susceptible lines.

To estimate the expected genetic gain across infection, we computed the mean genetic gain between first and second infection. We considered pedigree-based breeding values (eBV) as they were available across generations (geBVs were only available for G0 and G1 individuals) and were strongly correlated with geBV. eBVs were estimated from recorded phenotypes in G0, G1 and G2 individuals using a mixed model including fixed environmental effects and a random individual effect estimated from the pedigree relationship matrix (encompassing 1559 individuals) as implemented in the AIReml software (http://nce.ads.uga.edu/wiki/doku.php?id=readme.aireml). Genetic gain was expressed in genetic standard deviation (*σ*_g_) and was estimated within each line as the cross-product between the overall selection intensity (average of selection intensities in males and females) and the selection accuracy. Genetic gains from both lines were summed to yield the expected genetic divergence. Of note, the genetic gain in G2 was obtained by summing the gain from the mating between G1 sires and G1 dams on one hand, and that from the matings between G1 sires and G0 dams on the other hand.

To estimate putative trade-offs between resistance to GIN and lamb weight, measured body weights were modelled using a mixed model with repeated measures, including fixed effects (litter size, sex, generation, day post-infection, and an effect aggregating line and corresponding generation), and a random effect accounting for inter-individual variations. To account for differences in body weight at the beginning of the trial, weight data measured before any infection took place was fitted as a covariate.

### Host genotype x Environment (*G_h_ x E*) and Host genotype x Parasite (*G_h_ x G_p_*) interactions

To test for *G_h_ x E* and *G_h_ x G_p_* interactions, normalized FEC data collected at every time-point (24- or 30-day post-infection) were scaled within each experimental block (infection rank, day post-infection and considered environment) to prevent spurious signals from heterogeneous variances between blocks (Pollott & Greeff, 2004). A mixed model for repeated measures was built, whereby scaled normalized FEC were regressed upon two interaction terms (between the lamb genetic line and the time post-infection, or the lamb genetic line and the environment), and an additional random effect accounting for inter-individual variation.

Behaviour data were considered as normally distributed (Shapiro-Wilk’s test ranging between 0.91 and 0.98). Physical contact records were however skewed toward 0 and were thus binary encoded, *i.e*. 0 or 1 in absence or presence of contact with their mates or with the operator, and modelled using a logistic regression framework. To test for the effect of behaviour treatment, recorded behaviour data were regressed upon sheep line, sex and day and their interactions, and a random effect accounting for inter-individual variation. Regression models were built following stepwise variable selection procedure that aims to find the model with minimal Akaike Information Criterion (AIC). Bleating records in phase 1 of the test were also corrected for their initial value before chronic stress treatment took place to correct for the increased bleating in susceptible lambs from the stress group. Pearson’s correlations were estimated with the *rcorr()* function from the Hmisc package (Harrell & Dupont, 2017).

## Results

### Achieved divergence and response to selection

To evaluate the response to selection in both lines and our selection procedure accuracy, we compared the performance of both R and S sheep infected with *H. contortus* (Fig. 1, supplementary Fig. 1). G1 and G2 generations significantly diverged from the unselected G0 nucleus on the phenotypic scale: R lambs of respective generations showed FEC reduction of 0.62 (*t_70_* = −3.8, *P* = 2 x 10^−3^) and 0.67 σ_p_ (*t_20_* = −2.8, *P* = 5 x 10^−3^) relative to the G0 nucleus flock, and S lambs FEC deteriorated by 0.72 (*t_55_* = 4.99, *P* = 10^−6^) and 0.61 σ_p_ (*t_31_* = 3.11 *P* = 2 x 10^−3^) relative to their G0 unselected relatives (Fig. 1). This corresponded to phenotypic divergence between R and S lamb FEC across infection of 1.89 and 1.87 σ_p_ for G1 and G2 generations respectively (Fig. 1). Accordingly, 3.14 and 3.8 genetic standard deviations (σ_g_) were found between R and S lines at G1 and G2 generations (Fig 1), slightly higher than respective expected genetic gains of 2.13 and 2.66 σ_g_ expected genetic gains.

**Figure 1.**
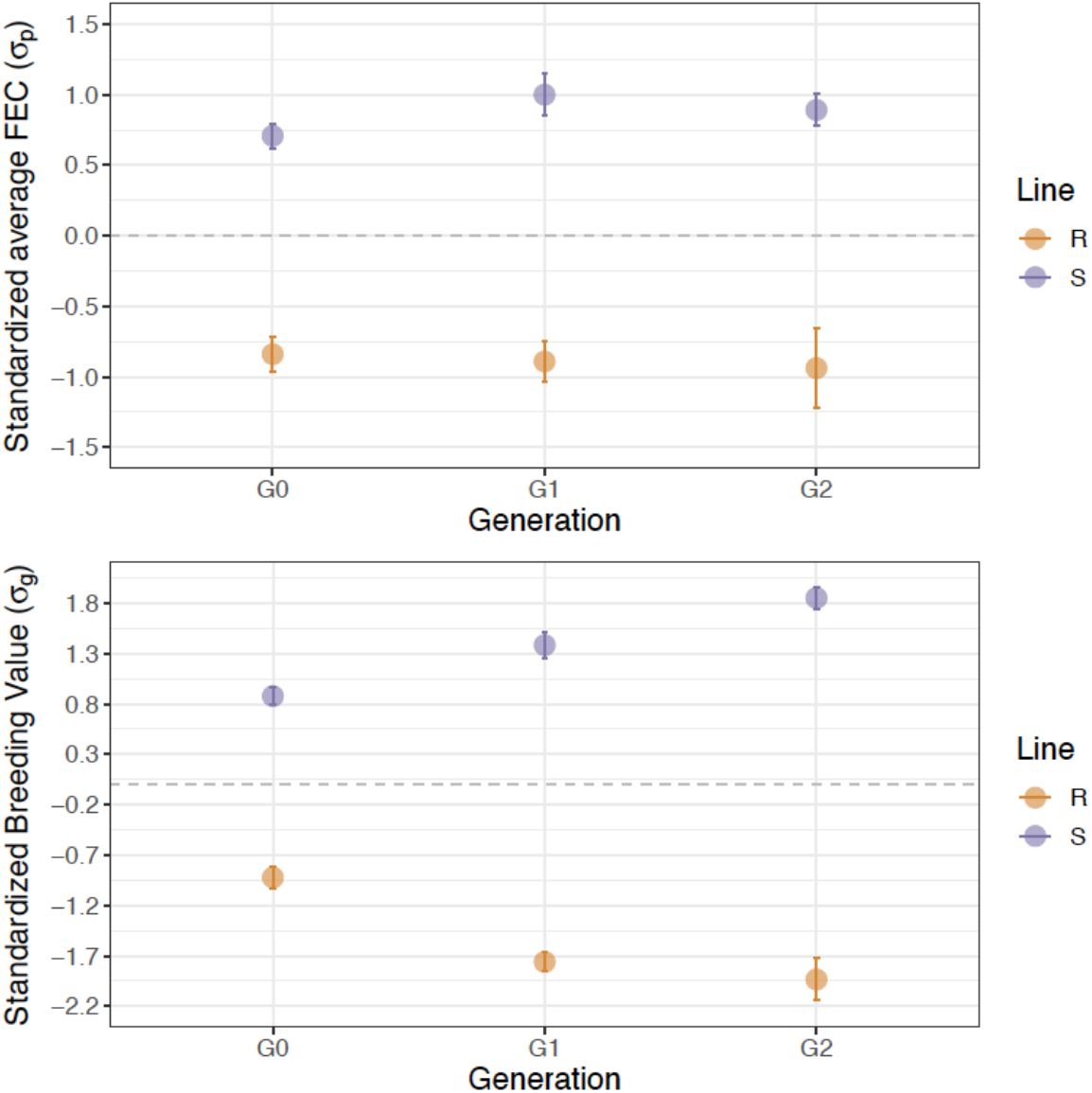
Achieved phenotypic and genetic divergences between sheep lines across generations. Top panels show the corrected average Faecal Egg Counts (mean ± s.e.) observed across two infections for parental (G0), first (G1) and second (G2) generations. Bottom panels represent the distribution of estimated breeding values (eBVs) using the pedigree information only. For the sake of comparison, FEC and eBVs data were scaled (mean centered and reduced to unselected G0 standard deviation unit). Grey dashed line indicates G0 mean value.

Despite similar selection intensity in R and S lines, observed response to selection for average FEC across infection was asymmetrical between lines in G2 lambs (Fig. 1). This asymmetry was evident from the regression of lambs response upon their midparent value (supplementary Fig. 1). In that case, the S line displayed a significant response to selection relative to their midparent value (supplementary Fig. 1, supplementary Table 2). This was accompanied by a reduced variance in FEC upon reinfection (supplementary Fig.1), with measured values at 30 dpi remaining high in a range between 1,650 and 14,250 eggs/g (median FEC = 5,625 eggs/g). On the contrary, the resistant line achieved half the response of their susceptible counterpart for FEC across infections (0.38 σ_p_ ± 0.27 and 0.82 σ_p_ ± 0.27 for the resistant and susceptible lines respectively, supplementary Table 2). This was the result of more variable FEC in that line (Fig. 1, supplementary Fig. 1), that was also evident at first and second infection (supplementary Fig. 1, supplementary Table 2). This lower response could not be related to inbreeding that was not significantly different between G2 R and S lines (*t_35_* = −2.01, *P* = 0.05).

In order to establish putative trade-offs between FEC and lamb growth, lambs were weighed. Analysis of their weight trajectories showed that they were not statistically different between selected lines (average weight differences of 568 g, *t_38_* = −0.61, *P* = 0.54 in G1 and of 1.27 Kg, *t_34_* = −1.31, *P* = 0.19 for G2 lambs), suggesting that higher resistance was not detrimental to production traits (supplementary Fig. 2, supplementary Table 2). However, the G2 R lambs were significantly lighter than their susceptible counterparts before any infection took place (4.3 Kg difference, F*_1,35_* = 8.99, *P* = 5 x 10^−3^). Of note, estimated geBVs and eBVs showed strong correlation (*r_241_* = 91% and 71% for FEC at first and second infection respectively, *P* < 10^−4^).

Altogether, we achieved significant divergence in GIN resistance between sheep lines with no evidence for detrimental effects on production traits.

### Sheep fully express their resistance potential under chronic stress but not after *T. colubriformis* infection

To identify putative G x E interactions, related individuals with divergent resistance to *H. contortus* infection were exposed to various environments, *i.e*. chronic stress or the intestinal parasite *T. colubriformis*. Half of the G1 selected lambs were exposed to chronic and unexpected stresses while the other half were maintained under controlled environmental conditions. Before chronic stress was applied, female lambs from the control group spent more time standing than female lambs in the other group (628 vs 578 min/day; F*_1,36_* = 6.47, *P* = 0.01, supplementary Table 2). This variable was hence not considered further. No significant difference was found in other behavioural data between both treatment groups (supplementary Table 2). Following exposure to chronic stress for five weeks and before any infection took place, lambs displayed altered behaviour. They bleated less in phase 2 of the arena test (2.35 count difference, F*_1,79_* = 6.09, *P* = 0.02, supplementary Table 2) and expressed less vigilance than their counterparts facing control conditions (0.69 count difference, F*_1,80_* = 5.06, *P* = 0.03, supplementary Table 2).

Despite significant alterations in sheep behaviour following chronic stress exposure, limited interactions were found between genetic line and their environment (Fig. 2; supplementary Fig. 3), as the relationship between genetic line and transformed FEC did not significantly differ across conditions (*F_1,82_* = 0.03, *P* = 0.87). However, phenotypic divergence between lines exposed to chronic stress significantly decreased 24 days after the second infection (Fig. 2). In that case, susceptible lambs excreted less eggs (−0.66 σ_p_, *t_252_* = −2.91, *P* = 4 x 10^−3^) than their sibs maintained under controlled conditions (Fig. 2).

**Figure 2.**
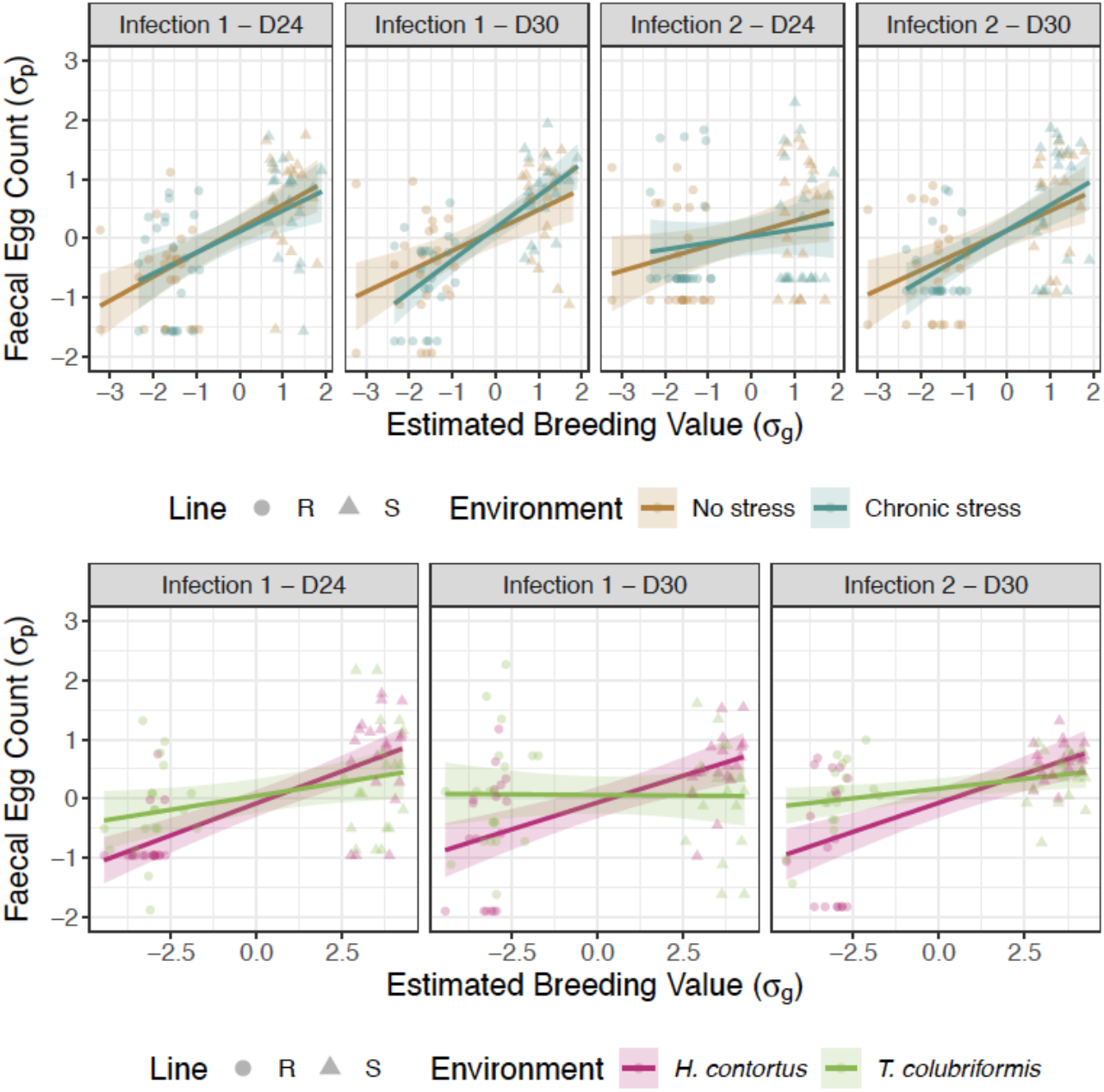
Genetic divergence stability across environments. The relationship between estimated breeding values (eBV) and Faecal Egg Count (FEC) across considered time points (infection rank - day post infection) and environments for resistant (circles) and susceptible (triangles) sheep lines. Top panels correspond to exposure to chronic stress and bottom panels illustrate the impact of infection by another parasite species. For the sake of comparison, raw FEC data were normalized with a 4^th^ root transformation and scaled (mean centered and reduced to standard deviation unit) within each group x time point, and eBVs were scaled within each trial.

The second trial aimed to investigate whether the genetic potential for resistance or susceptibility to *H. contortus* infection would be sustained in the face of another GIN species (Fig. 2). Of note, *T. colubriformis* infection yielded fewer eggs (average FEC of 411 eggs/g across conditions, ranging between 0 and 1,100 eggs/g) as a result of the lower fecundity of this parasite species. The phenotypic divergence between R and S sheep remained significant across considered GIN species (1.52*σ*_p_ difference, *F_1,73_* = 34.9, *P* = 3 x 10^−8^) but was largely driven by the existing divergence between *H. contortus* infected individuals. Indeed, the phenotypic expression of lamb genetic potential was significantly reduced after infection by the intestinal *T. colubriformis* species. In that case, lambs genetically susceptible to *H. contortus* were less affected by *T. colubriformis* infection, as evidenced by the mild difference in egg counts between both lines (138 eggs/g difference, *t*_36_ = 1.92, *P* = 0.06) and by the significant *G_h_ x G_p_* effect (−0.9 *σ*_p_, *F_1,73_* = 9.77, *P* = 0.003; Fig. 2).

The magnitude of *G_h_ x E* interactions were statistically different between sire families (supplementary Fig. 3): progenies of two susceptible sires displayed higher phenotypic variability following exposure to stressful conditions (supplementary Fig. 3). In some sire families chronic stress appeared to be beneficial (interaction term equal to −1.16 *σ*_p_, *P* = 0.024 for sire S-132550) while in others was detrimental (interaction term equal to 0.65 *σ*_p_, *P* = 0.046 for sire S-132361; supplementary Fig. 3). On the other hand, we found no evidence that the magnitude of *G_h_ x G_p_* interactions varied across families.

These two trials hence provide evidence that the phenotypic expression of *H. contortus* resistant individuals holds in the face of chronic stress, but can vary across families. It also offers a significant advantage in tolerance of infection by a different intestinal GIN species although to a lower extent than for the species used during selective breeding.

### Genomic and transcriptomic variations between *H. contortus* worms recovered in sheep lines

Another putative issue faced by the breeding of more resistant sheep, is the ability of parasites to adapt and circumvent the response of their host. To identify key parasite genes mediating survival of the host immune response, we compared the transcriptomic profiles of surviving *H. contortus* collected in both sheep backgrounds. On average 16 M (standard deviation of 5.66 M) reads were available for each library. One replicate collected from sheep 229 behaved as an outlier (supplementary Fig. 4). This difference was driving 38% of total variation in normalized RNA-seq read counts as shown in a principal components analysis (supplementary Fig. 4). This replicate was subsequently discarded from the dataset.

A limited set of ten genes appeared significantly differentially expressed (Benjamini-Hochberg adjusted *P* < 0.05) between worms from resistant and susceptible lambs (Table 1, Figure 3, supplementary Fig. 5). These ten genes exhibited significant reduction of their expression in worms from susceptible sheep with absolute fold-changes ranging between 2.3 and 16.7 (Table 1, Figure 3, supplementary Fig. 5). Accordingly, significant correlations were found between normalized transcript counts for these genes and matching sheep FEC values measured before necropsy (Table 1, supplementary Fig. 5): higher expression in worms was associated with lower FEC (*Pearson’s r* ranging between −0.74 and −0.87 across genes, *n* = 23). Among these ten genes (Table 1), seven were encoding cuticle collagen. This was either inferred from the function of their homologs in the model nematode *Caenorhabditis elegans (HCON_00007272, HCON_00007274, HCON_00182130*) or from domain prediction of their respective protein product (*HCON_00050580, HCON_00087240, HCON_00182130, HCON_00191700*, Table 1). Strongest differences and association with host phenotype were found for *HCON_00087240* and *HCON_00007274* that displayed expression levels 13.7 and 16.7 times higher in worms from resistant sheep.

**Table 1.** List of genes differentially expressed between *H. contortus* recovered from resistant and susceptible lines. Table lists the estimated fold changes and associated adjusted *P*-values for ten genes found differentially expressed in at least three pipelines. Their predicted genomic coordinates, gene features and orthology in *Caenorhabditis elegans* are given.

| Gene stable ID | RNAseq FC<sup>a</sup> | Adj. *P* | *r* with FEC<sup>b</sup> | qRT-PCR FC<sup>c</sup> | Chromosome | Coordinates (bp) | Gene feature<sup>d</sup> | *Caenorhabditis elegans* orthologs |
| --- | --- | --- | --- | --- | --- | --- | --- | --- |
| HCON_00029050 | 2.25 | 1.39E-03 | -0.80 | 1.29 | I | 40,762,263 - 40,774,522 | - | - |
| HCON_00191700 | 3.41 | 1.48E-02 | -0.86 | **2.72** | I | 4,916,615 - 4,917,957 | Collagen triple helix repeat (IPR008160); Nematode cuticle collagen, N-terminal (IPR002486) | - |
| HCON_00007272 | 4.92 | 4.50E-04 | -0.88 | **3.85** | I | 9,286,623 - 9,287,756 | - | col-176 |
| HCON_00007274 | 16.68 | 3.99E-03 | -0.79 | 1.77 | I | 9,295,404 - 9,296,508 | - | col-10, col-126, col-127, col-144, col-145 |
| HCON_00050580 | 2.50 | 1.39E-03 | -0.74 | **2.44** | II | 25,567,804 - 25,569,032 | Collagen triple helix repeat (IPR008160); Nematode cuticle collagen, N-terminal (IPR002486) | - |
| HCON_00087240 | 13.64 | 2.22E-02 | -0.75 | 0.89 | III | 31,491,433 - 31,496,242 | Nematode cuticle collagen and Collagen triple helix repeat domain containing protein | - |
| HCON_00090510 | 3.97 | 2.91E-03 | -0.74 | 4.41 | III | 36,604,400 - 36,605,347 | Secreted clade V protein (IPR035126) | - |
| HCON_00090580 | 2.62 | 1.48E-02 | -0.85 | 1.12 | III | 36,632,125 - 36,633,161 | Secreted clade V protein (IPR035126) | - |
| HCON_00132830 | 2.77 | 6.49E-04 | -0.87 | 1.12 | V | 2,187,669 - 2,196,836 | - | aqp-6 |
| HCON_00182130 | 3.12 | 3.50E-04 | -0.88 | **3.50** | X | 31,665,074 - 31,666,599 | Collagen protein | col-160, col-166, col-167, col-168, col-169, col-170, col-180 |

*a*: Fold-change (relative to worms from susceptible sheep). *b*: Pearson's *r* between gene-wise transcript count and Faecal Egg Count (*n* = 23, *P*-value ranged between 5.6 x 10<sup>-5</sup> and
4.15 x 10<sup>-8</sup>). *c*: Average qRT-PCR fold-change between resistant and susceptible worms collected from G2 lambs; bolded figures indicating significant differential expression after
Bonferroni correction (*P*<0.004). *d*: Interpro protein domains (IPRXXXXXX) were indicated for genes with no gene description available.

**Figure 3.**
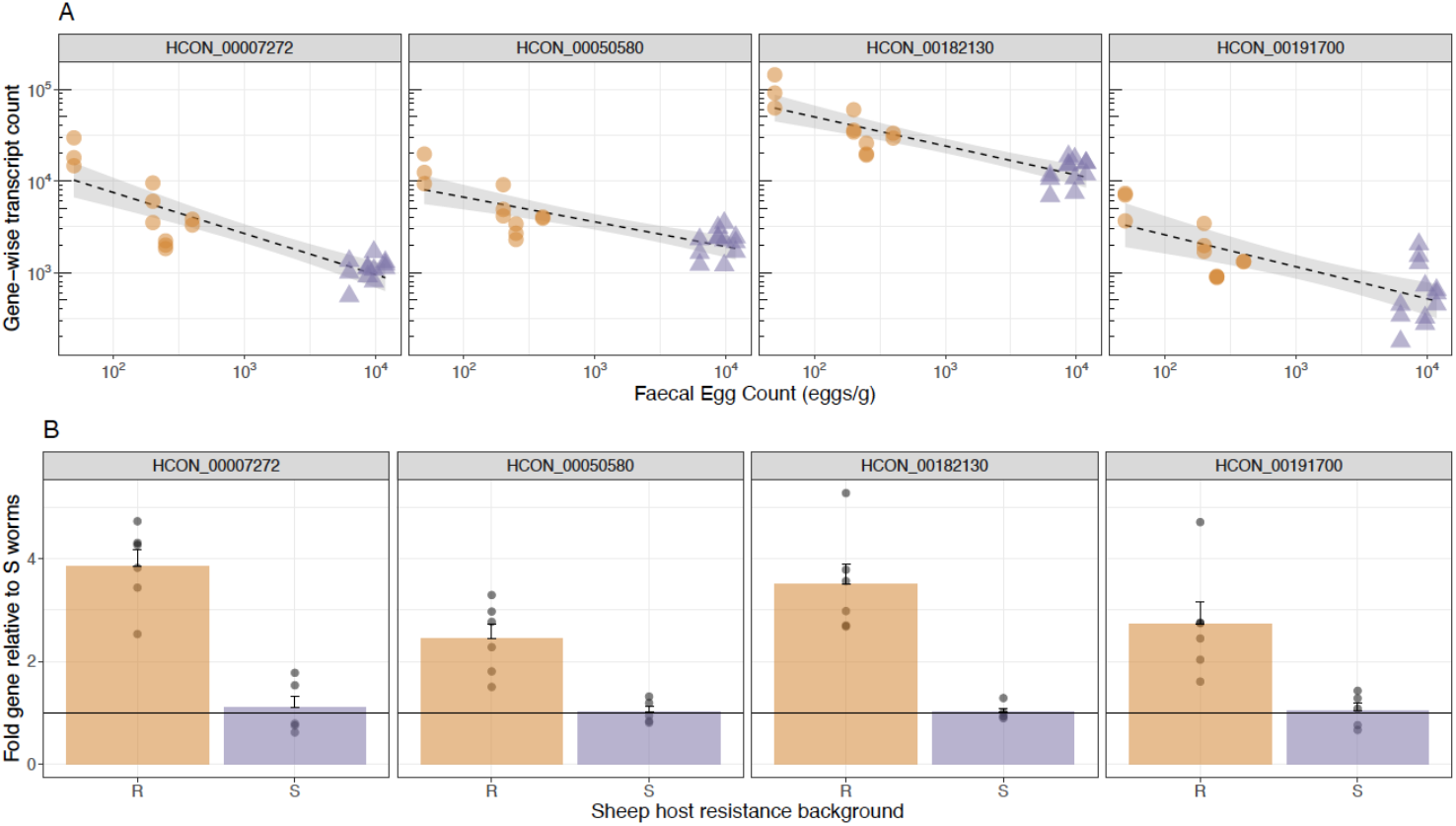
Expression profile of validated candidate genes associated with survival to resistant sheep in *Haemonchus contortus*. Top panels show the linear relationship between gene-wise transcript count estimated from RNAseq data in worms collected in 1st generation sheep and corresponding Faecal Egg Count (in eggs/g). Bottom panels give expression fold-change (relative to worms from susceptible sheep) estimated by qRT-PCR in worms collected from 2nd generation sheep. In that case, each dot stands for a biological replicate of 50 worms, and the horizontal line indicates null fold-change. Colors match sheep host background, resistant and susceptible in orange and purple respectively. *HCON_00007272*, a cuticular collagen homolog; *HCON_00050580*, a cuticular collagen domain containing protein; *HCON_00182130*, a collagen protein; *HCON_00191700*, a cuticular collagen domain containing protein.

Gene expression levels of nine of these genes were measured by qRT-PCR in an independent set of worms collected from G2 R and S sheep (Table 1, Figure 3). These data confirmed the increased expression for four cuticle collagen related genes in worm from R sheep (Figure 3), including *HCON_00007272* (fold change = 3.85 ± 0.78, *P* = 8.48 x 10^−9^), *HCON_00050580* (fold change = 2.44 ± 0.7, *P* = 8.69 x 10^−5^), *HCON_00182130* (fold change = 3.50 ± 0.99, *P* = 8.46 x 10^−8^), *HCON_00191700* (fold change = 2.72 ± 1.07, *P* = 1.94 x 10^−5^). Measured expression levels were not statistically different between conditions for other genes (Table 1).

We also evaluated how sheep resistance was constraining the genetic diversity of *H. contortus* populations by sequencing the DNA of pools of these worms from sensitive and resistant hosts. This strategy was applied to determine whether any major parasite gene would be involved in worm survival in resistant sheep. The genetic differentiation observed in pairwise comparisons involving worms from the same genetic line or from lambs with different genetic backgrounds was low (average F_ST_ of 0.018, ranging between 0.004 and 0.09; supplementary Fig. 6). This finding is compatible with low drift and lack of strong selection following a single infection. Nevertheless, we retrieved genes found within windows showing the most extreme levels of genetic differentiation between worms from different sheep backgrounds. Estimated genetic differentiation remained low over these windows (average F_ST_ of 0.033, ranging between 0.017 and 0.055) and encompassed a set of 37 genes found on every chromosome (supplementary Table 3). Among these, we found the *dpy-3* ortholog gene (*HCON_00165005*) is known to play a role in cuticle development in *C. elegans*, and might contribute to observed transcriptomic differences.

Altogether these findings suggest that an elevated transcription of cuticle component may concur to enhance survival of worms in resistant sheep, but the selection applied by the sheep genome is unlikely to select for a major gene in the parasite.

## Discussion

Understanding how the host-parasite system behaves following changes in their respective environments is central to ensure sustainable control of GIN in livestock through animal breeding. Our work investigated how directional selection for contrasting levels of resistance to GIN infection would affect expression of sheep potential toward environmental change. Because this selection induces abrupt shifts in the parasite niche we also established elements of the worm transcriptomic response across sheep backgrounds.

Our design relied on divergent sheep lines that provide a model system to quickly evaluate consequences of such environmental variation. We obtained aggregated estimates of the genotypic variance across environments, either quantified by the interaction between lamb genetic groups and their environment, or by sire reaction norms across environments. These observations suggest that environmental variations had limited effect on the phenotypic performances of genetically divergent sheep. But we found consistent increased expression of cuticle collagen components in the worms, compatible with an adaptive plastic response to their host immune response.

The response to selection was asymmetrical in G2 lambs. It yielded an increased divergence towards susceptibility rather than resistance, despite similar selection intensity across sexes and lines within generations. In the absence of replicated lines, it remains difficult to disentangle this observation from the differential contribution of random drift within each line (Falconer & Mackay, 1996). This may also indicate that the proportion of phenotypic variance of genetic origin is more difficult to estimate for resistant individuals than for susceptible lambs. Indeed, resistance is measured from FEC, whose distribution will be censored to 0 across a range of resistance levels, thereby hampering variance estimation. It is also possible that the selection applied to the pleiotropic gene networks underpinning the immune response to GIN infection (Lazzaro & Little, 2009; Sallé et al., 2012; Sparks et al., 2019) could yield asymmetric phenotypic expression upon selection. Additional rounds of divergence would be needed to support this hypothesis.

In line with previous reports (Gruner et al., 2004; Woolaston et al., 1990), susceptibility levels toward *H. contortus* infection were well correlated to that measured upon *T. colubriformis* infection and we found no indication of between sire variation across environments. But lambs selected for diverging susceptibility to *H. contortus* did not express the same divergence toward *T. colubriformis* infection. Previous estimated genetic correlations between FEC of both GIN species were positive but ranged between 0.9 and 0.76 at first and second infection (Gruner et al., 2004). This indicates that a large common genetic background is associated with resistance to both GIN species, but that a minor species-specific genetic component contributes to immune mechanisms associated with either the GIN species or with its niche (abomasum in one case or the small intestine in the other) or both. In contrast, only 38 genes were found differentially expressed across two divergent sheep lines bred for resistance to *H. contortus or T. colubriformis* and infected with *H. contortus* upon primary infection (Zhang et al., 2019). This intersecting set vanished following reinfection as differentially expressed genes were private to each sheep line (Zhang et al., 2019). These lines of evidence would suggest that despite a largely common genetic architecture between resistance to both GIN species, selection for resistance to one species may result in an efficient non-adaptive response at the transcriptomic level upon abrupt environmental modification (Ghalambor, McKay, Carroll, & Reznick, 2007). In our experiment, observed plasticity may also result from a similar maladaptive response originating from partially correlated traits in lambs selected for resistance to one species and exposed to another. This non-adaptive response may highlight cryptic mechanisms that have been maintained through time by selection, to provide selective advantage against seasonal variation in GIN community structures (O’Connor, Walkden-Brown, & Kahn, 2006). Selection would hence have acted on plasticity itself (Lynch & Walsh, 1998).

Of note, no significant differences in sheep phenotypes were found after chronic stress exposure apart from a reduced susceptibility of stressed lambs upon reinfection. The complex interactions between the immune response and chronic stress is primarily driven by the hypothalamic-pituitary-adrenal and the sympathetic-adrenal medullary axes, that respectively control the release of glucocorticoids and catecholamines (Khansari, Murgo, & Faith, 1990; Padgett & Glaser, 2003). The intricacies of both neuronal and immune system have not been fully resolved but evidence suggests that the glucocorticoid corticosterone dampens the immune response by promoting a shift from a pro-inflammatory Th-1/Th-17 response to a Th-2 response (Elenkov, 2004; Harpaz et al., 2013; Padgett & Glaser, 2003). This latter polarization is associated with beneficial outcome of GIN infection and enhanced in more resistant hosts, as reported in mice (Filbey et al., 2014) or in sheep (Terefe et al., 2007). The decrease in FEC observed in lambs exposed to chronic stress 24 days after reinfection is compatible with a delayed worm development that could be underpinned by an enhanced Th-2 response in these individuals. However, the lack of any beneficial effect at primary infection and the transient nature of this observation upon reinfection warrant further validation.

Directional selection of sheep with enhanced resistance potential to GIN may have consequences on parasite populations. Investigation of the genetic diversity of *H. contortus* pools of worms collected from the diverse sheep backgrounds revealed limited genetic differentiation. Our study was not designed to investigate the putative genetic adaptation of the worm population: resolving this question would request multiple rounds of artificial infection to evaluate the relative contributions of the random loss of alleles over time and that of the host background. Investigating micro-evolutionary trajectories in more resistant hosts would be faced with the difficulty of estimating allele frequencies in small census populations that could be partially overcome by appropriate controls, *e.g*. random sampling of small batches of individuals in the much bigger worm populations from susceptible sheep. It is striking however that genetic drift had a limited contribution in resistant sheep.

In contrast to the lack of differences at the genetic level, our observations revealed a consistent differential expression in a limited set of genes that mostly coded for cuticle collagen components. Nematodes are protected from environmental stressors by their cuticle, which is primarily composed of intricated collagen and collagen-like proteins (Page & Johnstone, 2007; Page, Stepek, Winter, & Pertab, 2014). Among the 122 known collagen-coding genes in *C. elegans*, 22 are associated with detrimental phenotypes when knocked-down (Page et al., 2014). However, the orthologs of none of these 22 genes were found to be differentially expressed in our set of genes. Instead, we found differential expression in *HCON_00007274* whose orthologs are known to be regulated by *elt-3 (col-144* ortholog) or upregulated (*col-126* and *col-127* orthologs) following *skn-1* and *dpy-7* RNA interference in *C. elegans* (Dodd et al., 2018). These authors also reported that the disruption of circumferential bands of the cuticle (annular furrows) could trigger distinct stress responses involving both *elt-3* and *skn-1* genes (Dodd et al., 2018). It may be speculated that worms from more resistant lambs suffered higher cuticle damage in relationship with a more effective response. This could select for worms with increased collagen turn-over to renew their cuticle. A controlled experiment exposing worms to eosinophil degranulation products from sheep with contrasted genetic status could help confirm this speculation. Alterations in GIN population plasticity may contribute to buffer *G_h_ x G_p_* interactions, thereby reducing the efficacy of selection imposed on the host population.

Our experimental trial found limited G x E interactions. This suggests that the selection of more resistant animals would hold across these conditions. However, the few interactions found suggest that individuals of higher genetic potential may have an increased environmental sensitivity. The selection of more resistant sheep was also associated with differential expression of *H. contortus* genes that may favour higher turn-over of cuticle components in worms recovered from resistant sheep.

Additional environmental disruption like feeding restriction could be investigated to confirm that the resistant potential holds under resource-limited environments and longer-term monitoring of the trajectories of both hosts and their parasites should establish the genetics of co-evolution between the two systems.

## Acknowledgements

This project was funded by the ANR Institut Carnot Santé Animale program (GEMANEMA project N° 11 CARN 016-01). GS received a Marie-Curie Agreenskills+ fellowship (grant agreement n°609398). NH and JAC were funded by the Wellcome Sanger Institute via their core funding of the Wellcome Sanger Institute (grant 206194).

## Data archiving

Raw reads produced in this project have been deposited at ENA with PRJEB23148 accession number. Genomic DNA sequence data is in accessions ERR2977490-ERR2977681, RNA-seq data is ERR3061846-ERR3061893. R scripts used for raw data analysis are freely available at https://github.com/guiSalle/GEMANEMA. Associated data files will be made available upon manuscript acceptance.

