## Supplementary Material for "Strongyle-resistant sheep express their potential across environments and leave limited scope for parasite plasticity"

**Supplementary Technical Note. Divergent sheep line creation**	**2**

**Diagram of the selection procedure**	**6**

**Supplementary Table 1. Primer pairs used for qRT-PCR and associated amplicon size and efficiency**	**7**

**Supplementary Table 2. Estimated contrasts and significance for considered fixed effects in implemented analyses**	**8**

**Supplementary Table 3. Genes found in genomic windows with outlying genetic differentiation between sheep lines**	**8**

**Supplementary Figure 1. Offspring to midparent Faecal Egg Count measured following H. contortus infection**	**10**

**Supplementary Figure 2. Weight trajectories measured in resistant and susceptible lambs following H. contortus infection**	**11**

**Supplementary Figure 3. Environmental effect on resistance potential varies across sire families**	**12**

**Supplementary Figure 4. Principal component analysis of RNAseq gene counts**	**13**

**Supplementary Figure 5. Relationship between lamb Faecal Egg Count and differentially expressed genes expression levels**	**14**

**Supplementary Figure 6. Genetic differentiation scan between pools of worms recovered from genetically susceptible or resistant sheep**	**15**

### Supplementary Technical Note. Divergent sheep line creation

In November 2013, 277 Romane lambs were experimentally challenged by two successive infections with *Haemonchus contortus*. Lambs received an ivermectin treatment 30 days after the first infection and a two-week recovery period was applied before the second infection took place. Faecal egg Count and Packed Cell Volume was measured just before the infection, 24, and 30 days post-infection. In total, 241 individuals were phenotyped on both occasions (Table 1).

Genotyping was performed for 274 of these lambs with a set of 1,000 SNPs, including markers in 8 QTL regions previously detected in a backcross between Romane and Black Belly breeds (Sallé et al., 2012). Sires (n = 37) were also genotyped for the 1,000 SNPs to assign missing sire-offspring kinship in 216 lambs. The total pedigree considered in the following analyses included 3 generations and 2,572 individuals.

**Table 1: Number of animal by sex and Faecal Egg Count measures among the 274-candidate animals**

|  | males | females | total |
| --- | --- | --- | --- |
| Phenotyped at 1st infection | 133 | 122 | 255 |
| Phenotyped at 2nd infection | 132 | 125 | 257 |
| Phenotyped in both infections | 121 | 120 | 241 |

The founder individuals mated to create the first generation of divergent individuals (G1) were selected among 274 animals with phenotype and genotype information. Their breeding value was estimated following a single-step Best Linear Unbiased Prediction (BLUP) approach that uses both the genomic and pedigree information (with equal weights).

**Table 2: List of sires to create generation G1**

| Selected sire | Sire Status | Mother | Father | sire gBV | Number of G1 offsprings |
| --- | --- | --- | --- | --- | --- |
| 20000132336 | R | 20000120766 | 20000120424 | -2.10 | 51 |
| 20000132471 | R | 20000120075 | 20000120140 | -1.21 | 39 |
| 20000132453 | R | 20000121413 | 0 | -1.20 | 46 |
| 20000132361 | S | 20000120399 | 20000120375 | 1.07 | 41 |
| 20000132497 | S | 20000121045 | 20000120985 | 1.18 | 44 |
| 20000132550 | S | 20000120415 | 20000120985 | 1.30 | 13 |

Individuals with most extreme breeding values at first and second infection were selected, yielding 3 sires at each extreme end of the phenotypic range (Table 2). Of note, two of the susceptible sires were half-sibs hence reducing the sampling effort (lower *Ne*) of available genetic variability in the susceptible lineage (Falconer and Mackay, 1996). In ewes, the selection intensity was lower, *i.e.* the median breeding values was chosen as a cut-off, to ensure enough G1 lambs would be produced.

In March 2015, 236 G1 lambs were born (Table 3), out of which 80 individuals were needed for experimental groups. Among the 236 born lambs, 180 individuals were selected according to their expected breeding values (average of their parents genomic breeding values across 1st and 2nd infection) for genotyping using the same 1000-SNP panel. Their genomic breeding values were estimated using the G0 founder population as a reference panel. Based on the estimated genetic merit, 91 G1 individuals (43 males & 48 females) were selected to ensure a minimum difference of 0.5 standard deviation between the genomic breeding values of both lines and 1 standard deviation between their pedigree-based breeding values. For the first experimental trial (summer 2015), 87 individual lambs were ultimately enrolled to measure genotype x environment interactions mediated by chronic stress.

**Table 3. number of animal by sex, line and stress conditions**

|  | R males | R females | S males | S females | Total |
| --- | --- | --- | --- | --- | --- |
| Chronic stress | 10 | 13 | 11 | 10 | 44 |
| Enrichment | 11 | 12 | 10 | 10 | 43 |
| Total | 21 | 25 | 21 | 20 | 87 |

A second generation of lambs was subsequently created (G2). The most extreme G1 lambs were chosen to retain 3 sires within each line (Table 4). To increase the genetic gain in females, 82 females were selected from both G0 and G1 ewes, ensuring 2 standard deviations between the average expected breeding values of the two lines. Controlled matings were performed between individuals with limited kinship ensuring an expected inbreeding coefficient in the G2 lambs lower than 0.04. Lambing took place in 55 out of the 82 selected ewes and generated 111 offsprings, 80 of which were selected according to the expected breeding values (average of their parents genetic merits ; Table 5).

**Table 4. List of sires used to create the second generation G2**

| Sire of G2 | Grand Sires of G2 | status |
| --- | --- | --- |
| 2000152347 | 20000132336 | R |
| 2000152365 | 20000132471 | R |
| 2000152226 | 20000132453 | R |
| 2000152258 | 20000132361 | S |
| 2000152337 | 20000132497 | S |
| 2000152434 | 20000132550 | S |

**Table 5: Number of males and females by lines**

| Line | males | females | total |
| --- | --- | --- | --- |
| R | 31 | 23 | 54 |
| S | 26 | 31 | 57 |
| total | 57 | 54 | 111 |

### Diagram of the selection procedure

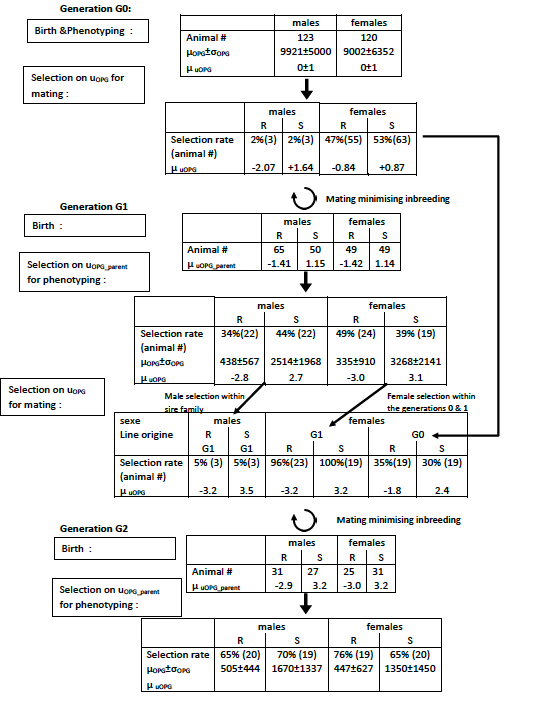

### Supplementary Table 1. Primer pairs used for qRT-PCR and associated amplicon size and efficiency

| **Gene** | **Forward primer** | **Reverse primer** | **Amplicon size (bp)** | **Efficiency (%)** |
| --- | --- | --- | --- | --- |
| *ama* | TATGGGAGGTCGTGAAGGTC | GTGGGCTTCATAGTGGGCATA | 214 | 104 |
| *far* | TGCCAAGGACTATGCCAAGT | TGAGTGCGTCGATCTTTCCC | 128 | 100 |
| *gpd* | ACGAGACCTACAATGCAGCC | GCGAGACAGTTGGTGGTACA | 67 | 101 |
| *HCON_00007272* | CTTCCACCATGGTGCGTCTG | GCCAGGGTTACCAGGTTGTC | 95 | 106 |
| *HCON_00007274* | CCATGGTGTGTTTGCGAGGT | AGGGTTACCAGGTTGTCCAGG | 86 | 101 |
| *HCON_00029050* | CAACTCATTCTGCTCCTCATTTGC | ACCTCCTCCTCCGAAGCAAG | 71 | 104 |
| *HCON_00050580* | GGGGCGAGTATACCAGAATCCA | CAGCATGACGATACCTGCCG | 66 | 107 |
| *HCON_00087240* | TGGCGCGTCTCTGATTATCG | CATCGGTAGCAACCCGGAAA | 118 | 95 |
| *HCON_00090510* | CCTACAACCGTGACCTGGTC | ATCTGCGAAAGGGCTATCGG | 119 | 115 |
| *HCON_00090580* | GGTGTGGACTGTGATAAGCTGAG | TGTCGACAGCCCTTTTGCAG | 56 | 102 |
| *HCON_00132830* | TATGGCAGCTGGCGCTATCT | CCAAACGATCGTGCAGGGTT | 53 | 107 |
| *HCON_00182130* | CACAACAACGATGAGCAACGAC | ACCGAAAAAGGCGACTCTGC | 93 | 108 |
| *HCON_00191700* | CAAGGCGGAGAATGGGCTTA | TGAGACTGAAGGTGGTGGCA | 77 | 100 |

### Supplementary Table 2. Estimated contrasts and significance for considered fixed effects in implemented analyses

See attached file.

Table provides output summary for every reported modeling of response to selection, asymmetry of that response, weight trajectory, behaviour data, genotype x environment interaction across or within sire families. For every summary, contrasts between reference and other factor levels are given with respective standard errors (s.e.), t-test values and *P*-value. Any significant difference (*P* < 0.05) is highlighted in green.

### Supplementary Table 3. Genes found in genomic windows with outlying genetic differentiation between sheep lines

| **Gene** | **Chromosome** | **Gene start (bp)** | **Gene end (bp)** | ***Caenorhabditis elegans* ortholog** |
| --- | --- | --- | --- | --- |
| HCON_00022360 | I | 31480931 | 31491392 |  |
| HCON_00025640 | I | 36377419 | 36381726 | *C45B11.6* |
| HCON_00036120 | II | 3581831 | 3582337 |  |
| HCON_00049130 | II | 22477177 | 22494299 | *Y25C1A.13* |
| HCON_00049880 | II | 24424834 | 24425196 |  |
| HCON_00193140 | II | 29030456 | 29047041 |  |
| HCON_00052990 | II | 29044594 | 29045188 |  |
| HCON_00091320 | III | 37747395 | 37748133 |  |
| HCON_00094050 | III | 41471354 | 41481092 |  |
| HCON_00094060 | III | 41486869 | 41501504 | *sbp-1* |
| HCON_00109020 | IV | 16989747 | 17012821 | *scav-4* |
| HCON_00121240 | IV | 36752725 | 36764291 |  |
| HCON_00128300 | IV | 47832401 | 47838470 |  |
| HCON_00136900 | V | 7876408 | 7882212 |  |
| HCON_00140940 | V | 14033038 | 14034046 |  |
| HCON_00141740 | V | 15377695 | 15385120 |  |
| HCON_00143410 | V | 17943640 | 17949075 |  |
| HCON_00143420 | V | 17949129 | 17951366 |  |
| HCON_00146000 | V | 22919855 | 22921077 |  |
| HCON_00164360 | X | 935195 | 937008 |  |
| HCON_00165005 | X | 1909134 | 1910371 | *dpy-3* |
| HCON_00165250 | X | 2274629 | 2289564 | *rgl-1* |
| HCON_00166090 | X | 3528143 | 3530792 | *R09F10.3* |
| HCON_00168010 | X | 6864144 | 6887745 | *ldb-1* |
| HCON_00168080 | X | 7064091 | 7073013 | *idhb-1* |
| HCON_00168280 | X | 7357794 | 7364054 | *wdr-5.2* |
| HCON_00169420 | X | 9371799 | 9379947 | *Y8A9A.2* |
| HCON_00169480 | X | 9408326 | 9427816 |  |
| HCON_00184620 | X | 35549878 | 35554073 |  |
| HCON_00185600 | X | 36966654 | 36972546 |  |
| HCON_00185700 | X | 37143278 | 37160937 | *mlck-1* |
| HCON_00186210 | X | 37823153 | 37833034 | *exc-1* |
| HCON_00186770 | X | 38787409 | 38799293 |  |
| HCON_00188290 | X | 40861120 | 40888652 | *T23G5.2* |
| HCON_00188540 | X | 41259509 | 41265941 | *kynu-1* |
| HCON_00190240 | X | 43879597 | 43880192 |  |
| HCON_00190250 | X | 43880233 | 43884428 |  |

### Supplementary Figure 1. Offspring to midparent Faecal Egg Count measured following *H. contortus* infection

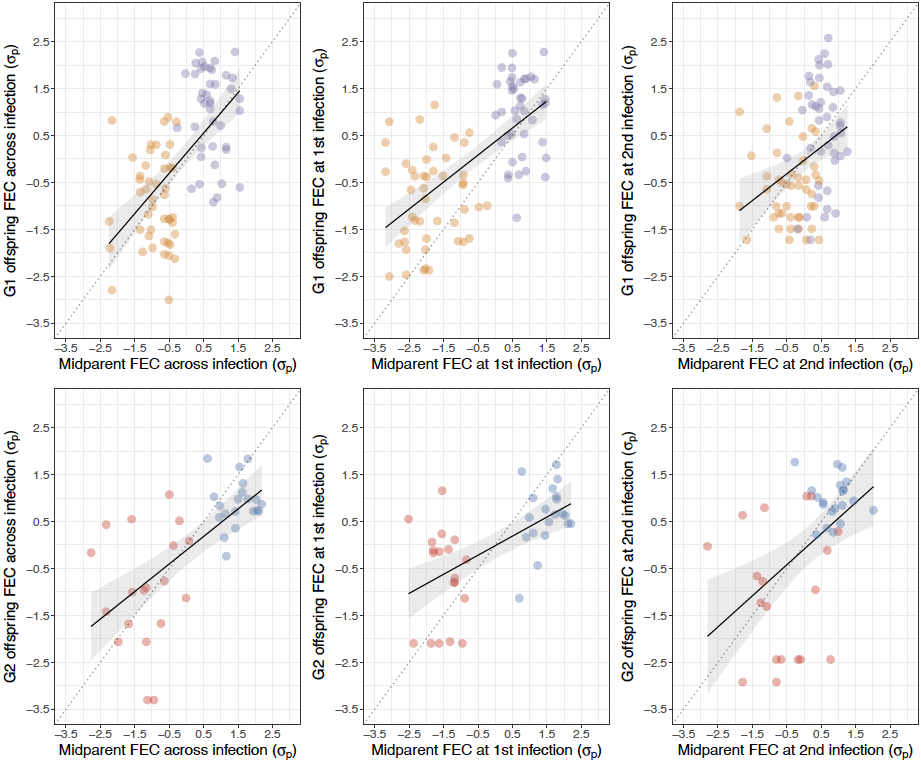

The figure represents average Faecal Egg Count (FEC) of *H. contortus* infected lambs, plotted against their respective midparent values (mean phenotype values of respective sire and dam). Top and bottom panels correspond to data gathered in the first (G1) and second (G2) generation of divergence respectively, with dot colors matching genetic lines (orange and red for resistant individuals in G1 and G2 respectively). For each infection, FEC were averaged across two measures recorded at 24 and 30 days after 1st (middle) or 2nd (right) infection or across infections (left), normalized (fourth-root transformed) and corrected for environmental effects. Resulting corrected FEC were then standardized (mean centered and reduced to G0 standard deviation unit, 𝜎_p_) for the sake of comparison across generations. Slope of the regression curve (black line with 95% confidence interval in gray) approximates realized trait heritability and dotted line materializes complete genetic response (heritability of 1).

### Supplementary Figure 2. Weight trajectories measured in resistant and susceptible lambs following *H. contortus* infection

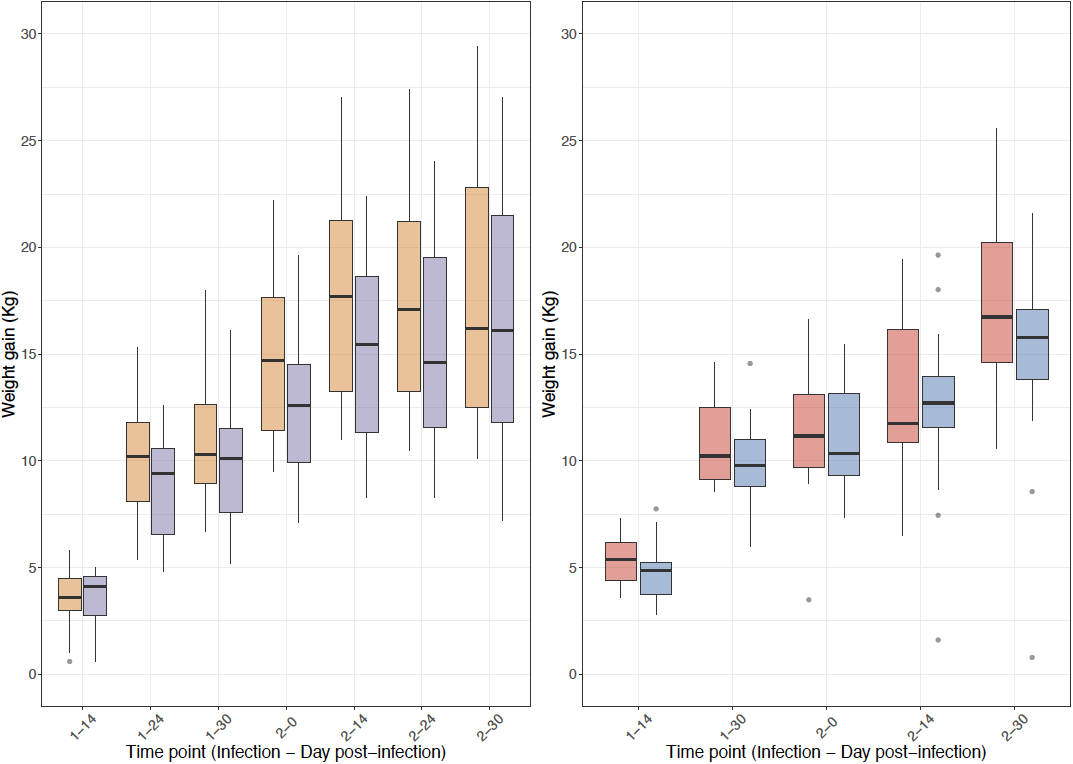

Box-plot of measured body weight data for resistant (orange or red) and susceptible (purple or blue) sheep lines. Phenotypes recorded on first generation lambs (G1) are given in top panels, and bottom panels present that measured on second generation lambs (G2). Horizontal bar stands for the median value within each sheep line and time point.

### Supplementary Figure 3. Environmental effect on resistance potential varies across sire families

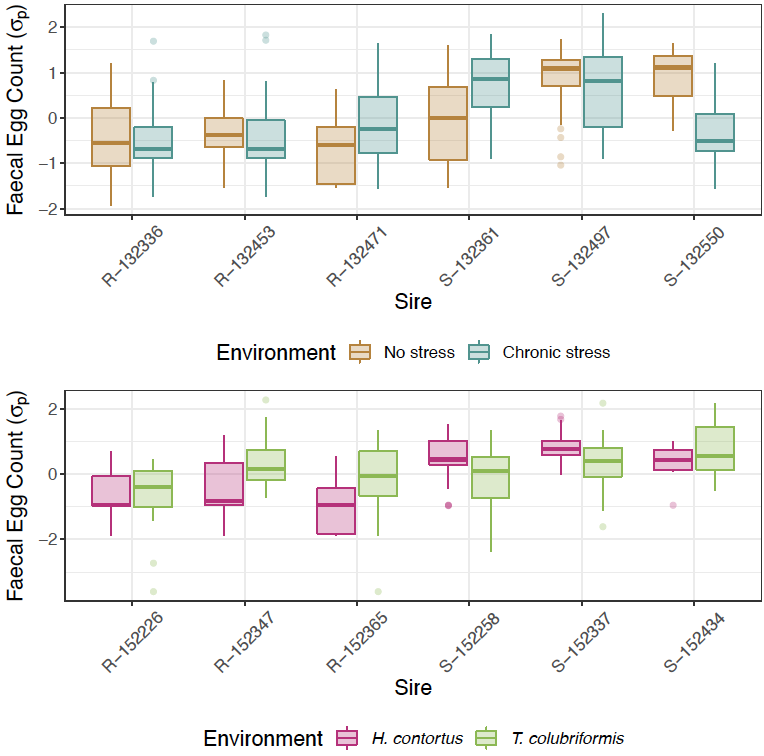

This figure represents the within-sire family variation in Faecal Egg Count (FEC, given in phenotypic standard deviation 𝜎_p_) across environmental conditions. FEC were normalized (fourth-root transformation) and then mean centered and scaled to unit standard deviation within each experimental block for the sake of comparison across conditions. First sire name letter indicates the divergent line it belongs to (R, resistant or S, susceptible). The picture indicates that the magnitude of G x E interaction varies among sires across stress conditions (top panel): sires S-132361 and S-132550 had their progenies showing higher or lower susceptibility under chronic stress respectively. The trend in resistance was conserved across GIN species in every sire family (bottom panel).

### Supplementary Figure 4. Principal component analysis of RNAseq gene counts

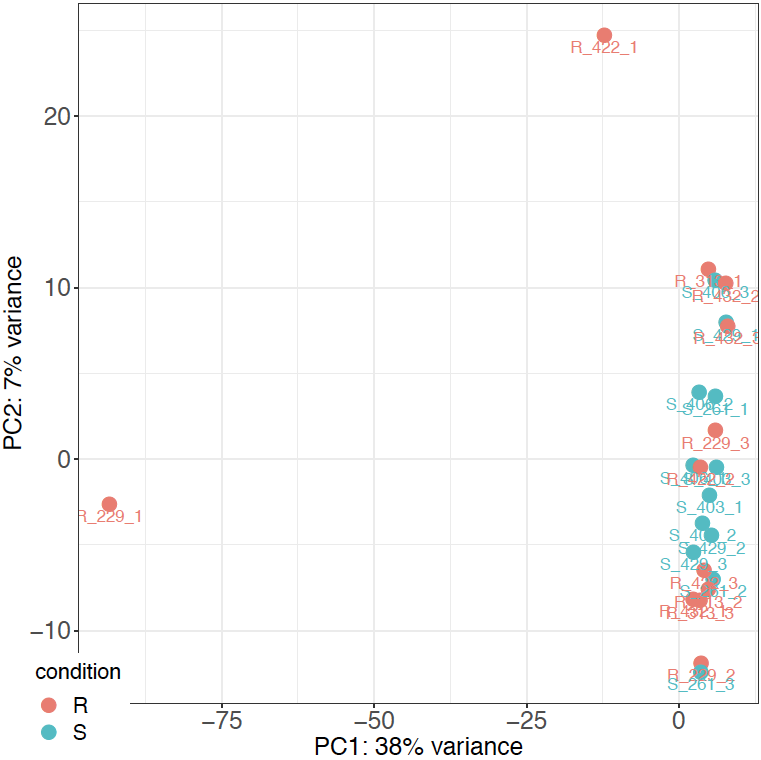

The figure represents the outlying contribution of one replicate pool of worms collected from sheep 229 to the total variance in gene transcription level.

### Supplementary Figure 5. Relationship between lamb Faecal Egg Count and differentially expressed genes expression levels

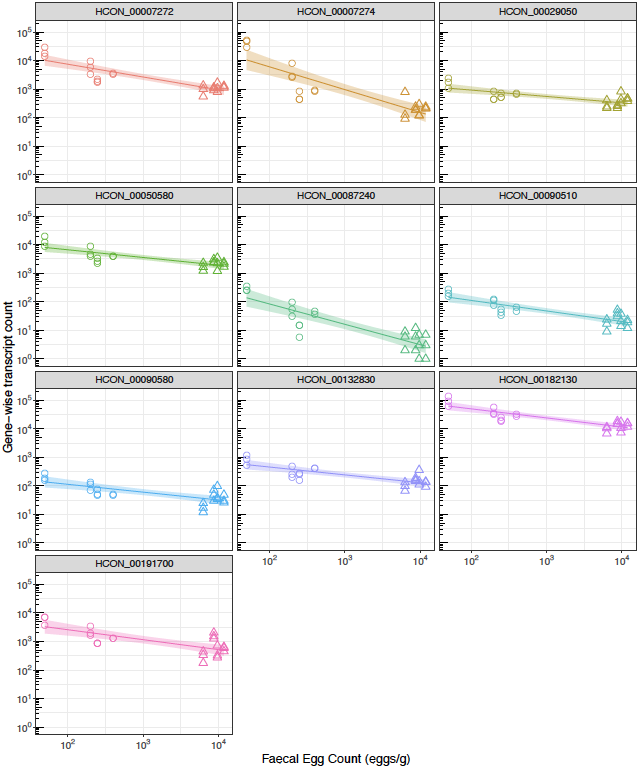

This figure shows the linear relationship between Faecal Egg Count measured 30 days after re-infection (in eggs/g) and gene expression levels measured by gene-wise transcript counts obtained with the STAR aligner in worms collected from resistant (circle) and susceptible (triangle) lambs. ¶

### Supplementary Figure 6. Genetic differentiation scan between pools of worms recovered from genetically susceptible or resistant sheep

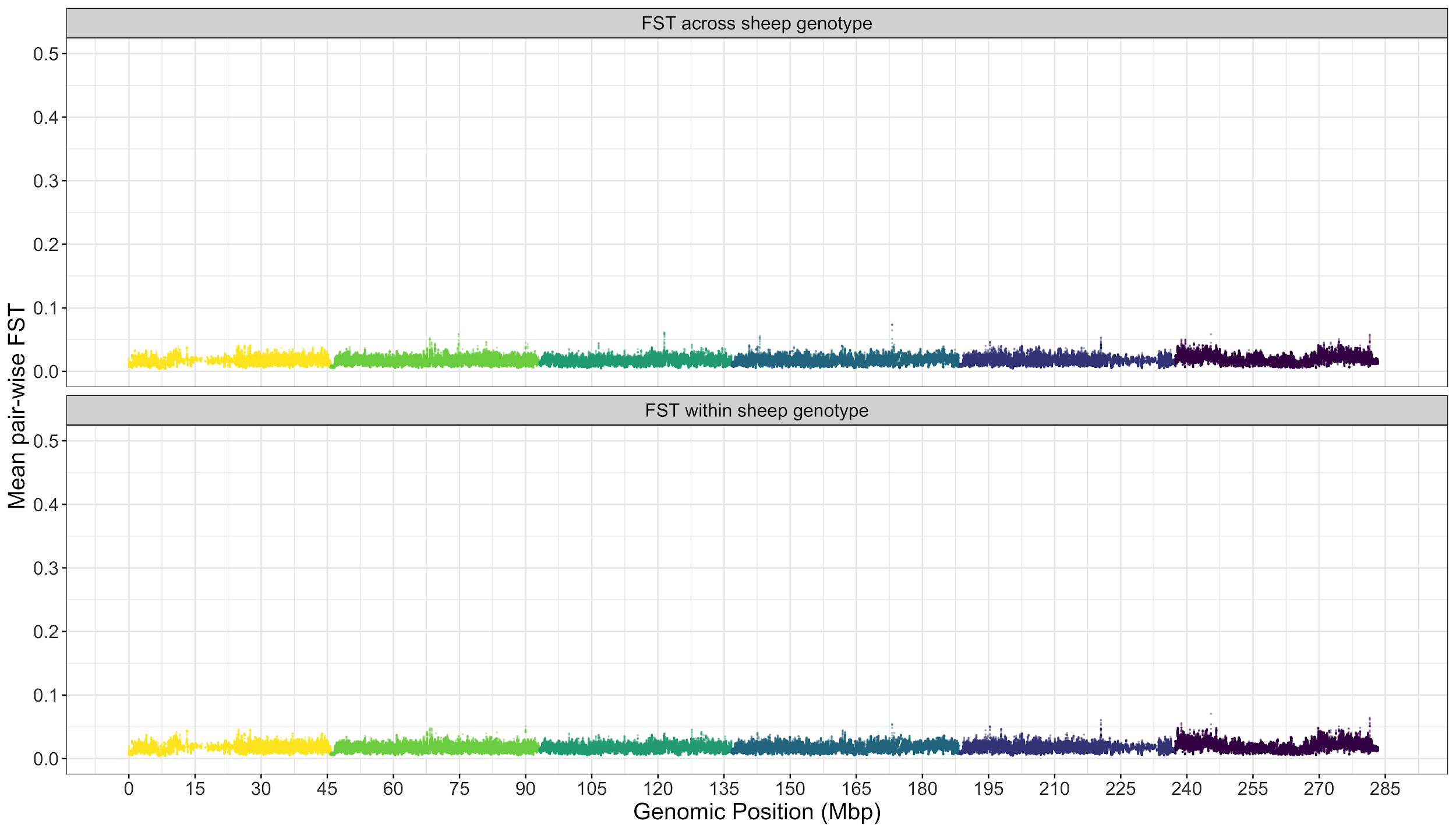

Average pairwise F_ST_ coefficients estimated between worm populations recovered from different (FST across sheep genotype) or same (FST within sheep genotype) is plotted against genomic position. Colors refer to the five autosomes (ranging from chromosome I in yellow to V in blue), and the sex chromosome in dark purple.
